# The need to include phylogeny in trait-based analyses of community composition

**DOI:** 10.1101/084178

**Authors:** Daijiang Li, Anthony R Ives

**Affiliations:** Department of Botany, University of Wisconsin-Madison, Madison, WI 53706, USA; Department of Zoology, University of Wisconsin-Madison, Madison, WI 53706, USA

**Author notes:** Present address: Department of Wildlife Ecology and Conservation, University of Florida, Gainesville, FL 32611, USA. Author contributions: DL and AI designed the study; DL conducted the simulations; DL and AI wrote the manuscript.

**Keywords:** Phylogeny, functional traits, fourth-corner problem, trait-environment relationship, phylogenetic linear mixed model

## Abstract

1. A growing number of studies incorporate functional trait information to analyse patterns and processes of community assembly. These studies of trait-environment relationships generally ignore phylogenetic relationships among species. When functional traits and the residual variation in species distributions among communities have phylogenetic signal, however, analyses ignoring phylogenetic relationships can decrease estimation accuracy and power, inflate type I error rates, and lead to potentially false conclusions.
2. Using simulations, we compared estimation accuracy, statistical power, and type I error rates of linear mixed models (LMM) and phylogenetic linear mixed models (PLMM) designed to test for trait-environment interactions in the distribution of species abundances among sites. We considered the consequences of both phylogenetic signal in traits and phylogenetic signal in the residual variation of species distributions generated by an unmeasured (latent) trait with phylogenetic signal.
3. When there was phylogenetic signal in the residual variation of species among sites, PLMM provided better estimates (closer to the true value) and greater statistical power for testing whether the trait-environment interaction regression coefficient differed from zero. LMM had unacceptably high type I error rates when there was phylogenetic signal in both traits and the residual variation in species distributions. When there was no phylogenetic signal in the residual variation in species distributions, LMM and PLMM had similar performances.
4. LMMs that ignore phylogenetic relationships can lead to poor statistical tests of trait-environment relationships when there is phylogenetic signal in the residual variation of species distributions among sites, such as caused by unmeasured traits. Therefore, phylogenies and PLMMs should be used when studying how functional traits affect species abundances among communities in response to environmental gradients.

## Introduction

Species composition and abundance in ecological communities depend in part on both the environmental conditions at a site and the traits expressed by species that allow them to live under these environmental conditions. Typically, environmental conditions at a site allow only a subset of species from the regional species pool to reach high abundances, with different functional traits favouring species in different sites. Therefore, both environmental conditions and functional traits play an important role in explaining species abundances in communities. To better understand community assembly, we need to study the statistical interaction between environmental conditions at a site and the functional traits of species that live there (McGill et al. 2006; Westoby & Wright, 2006).

Common statistical approaches to analyse how traits mediate species responses to environmental variables have used either ordination with permutation tests (the fourth-corner problem and RLQ analysis, Legendre, Galzin & Harmelin-Vivien 1997; Dray & Legendre, 2008) or an indirect two-step approach. The fourth-corner problem links three data matrix tables: a site × species incidence/abundance matrix (L), a site × environmental variables matrix (R), and a species × traits matrix (Q). The traits × environmental variables matrix (R’LQ) is the fourth matrix (thus explaining the etymology of the approach). While this approach provides a good qualitative overview of how traits and environmental variables are associated, it does not give information about species-specific variation in responses to environmental variables, and it is difficult to use for prediction. The second, two-step approach first fits species-specific regressions of abundance against environmental variables; the resulting regression coefficients are then regressed against traits (e.g., Soudzilovskaia et al. 2013). This approach, while informative at the species level, does not incorporate all community data in a single analysis and has low statistical power (Jamil et al. 2013).

The interactions between traits and environmental variables can also be directly tested with model-based methods (Bolker et al. 2009; Jamil et al. 2013; Brown et al. 2014; Warton et al. 2014, Ovaskaine, De Knegt & Delgado 2016). Statistically, the interaction between traits and environmental variables can be estimated as the trait-environment interaction coefficient in generalized linear models (GLMs, Brown et al. 2014), linear mixed models (LMMs, Ovaskaine, De Knegt & Delgado 2016), or generalized linear mixed models (GLMMs, Pollock, Morris & Vesk 2012; Jamil et al. 2013). These model-based methods allow model selection and prediction, and are often more flexible, powerful, and informative than fourth-corner and two-step approaches (Ives & Helmus 2011; Jackson et al. 2012; Brown et al. 2014; Warton et al. 2014).

Most analyses of trait-environment interactions ignore phylogenetic relationships among species, despite the large literature on phylogenetic analyses in comparative studies (Felsenstein 1985; Harvey & Pagel 1991; Paradis 2012; Garamszegi 2014) and the relevance of phylogeny to many areas of ecology (Webb et al. 2002; Cavender-Bares et al. 2009). This can lead to statistical problems because functional traits often exhibit a phylogenetic pattern in which closely related species share similar trait values (i.e., phylogenetic signal, Blomberg, Garland & Ives 2003). If there are multiple traits that affect species abundance or incidence (or other characteristic of interest), then the unmeasured traits with phylogenetic signal may generate covariance in the unexplained, residual variation after accounting for measured traits. This covariance in the residual variation will reflect the phylogeny, and will affect model estimation and hypothesis testing of regression coefficients (e.g., Felsenstein 1985, Martins & Hansen 1997; Garland, Bennett & Rezende 2005; Revell 2010).

Here, we investigate the need to incorporate phylogenetic covariance among species in regressions for trait-environment interactions. We considered a regression problem in which there is a causal but unmeasured (latent) trait that introduces unexplained variability in species abundance, and phylogenetic covariance in the unexplained variation if the unmeasured trait has phylogenetic signal. This gives four possible cases (Revell 2010): the pairwise combinations of whether or not there is phylogenetic signal in the measured trait in the regression, and whether or not there is phylogenetic signal in the residual variation. We then compared the accuracy, type I error rates, and statistical power of linear mixed models (LMMs) and phylogenetic linear mixed models (PLMMs, Ives & Helmus 2011) in estimating the trait-environment interaction coefficient. We show that when there is phylogenetic signal in the residual variation (latent trait), PLMM outperformed LMM, with LMM performing particularly poorly when there is also phylogenetic signal in the measured trait.

## Materials and methods

We simulated data to test the importance of accounting for phylogenetic relationships when studying how functional traits interact with environmental variables to affect species abundances. All simulations and calculations were performed with R (R Core Team, 2015).

### Simulations

We simulated the abundance *Y* of species *j* (*j* = 1,…, *n*) at site *s* (*s* = 1,…, *m*) that depends on two site environmental variables (env1 and env2) and two species functional traits (trait1 and trait2) using the model

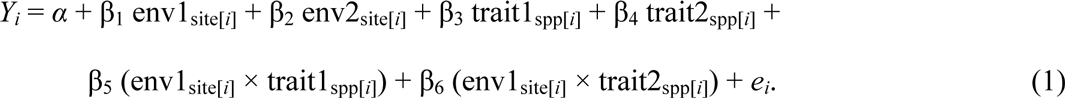

Functions spp[*i*] and site[*i*] map the observation *i* to the identity of the species and site, respectively (Gelman & Hill, 2007, p251–252), so *i* takes values from 1 to *nm*. We assume both environmental variable env1 and functional trait trait1 are measured. Env1 (e.g., soil fertility, canopy cover) affects the abundance of all species among sites (β_1_ ≠ 0), and trait1 (e.g., nutrient absorption capacity, specific leaf area) determines in part the overall abundance of species (β_3_ ≠ 0). Furthermore, there is an interaction between env1 and trait1 (β_5_ ≠ 0) implying that trait1 affects the performance of species along the environmental gradient env1.

To introduce unexplained variation and phylogenetic signal, we treated env2 and trait2 as unmeasured (latent) variables. Like env1, env2 has a direct effect on species abundances (β_2_ ≠ 0). Like trait1, trait2 determines in part species abundances (β_4_ ≠ 0) and has an interactive effect with env1 (β_6_ ≠ 0). As we are mainly interested in the trait × environment interactions for the measured data (env1 and trait1), we did not include the interactions between env2 and trait1 or trait2. Our goal is to investigate the interaction between env1 and trait1 which is given by β_5_. Consequently, we set all parameters in equation 1 other than β_5_ to be 1. Finally, we simulated *e_i_* as a normal random variable that is independent among species and sites. In this way, we treated the abundance of species *Y* as log-transformed values from count data. We did not simulate abundance as raw count data because log-transformation of count data usually does not affect the significance tests for regression coefficients when low count values (<5) are uncommon (Ives 2015; Warton et al. 2016).

We simulated the phylogeny as a uniform birth-death process with birth rate = 1 and death rate = 0 using the sim.bdtree function of the geiger R package (Harmon et al. 2008). The phylogeny gives the expected phylogenetic covariances among species under Brownian motion evolution (Grafen 1989; Martins & Hansen 1997) that can be used to construct a matrix **C**, and when there is no phylogenetic signal the (zero) covariance structure is given by the identity matrix **I**. Because functional traits may or may not have phylogenetic signal, we simulated four scenarios for the two functional traits: trait1 with phylogenetic signal but not trait2 (trait1: **C**; trait2: **I**); trait2 with phylogenetic signal but not trait1 (trait1: **I**; trait2: **C**); both traits with phylogenetic signal (trait1: **C**; trait2: **C**); and neither trait with phylogenetic signal (trait1: **I**; trait2: **I**). Functional traits without phylogenetic signal were simulated as ***N***(0, 1) normal random variables; functional traits with phylogenetic signal were simulated using the fastBM function of the phytools R package (Revell 2012). We simulated env1 as a uniform distribution ranging from –1 and 1 to generate a strong environmental gradient. Variable env2 and residuals *e*_i_ were simulated as ***N***(0, 1) normal random variables.

We conducted simulations with 30 sites. To study type I error rates (false positives that incorrectly reject the true null hypothesis), we set β_5_ = 0 and varied the number of species (20, 30, 40, 50, 60, 70, 80). To study statistical power, we varied the value of β_5_ (0, 0.25, 0.5, 0.75, 1) and fixed the number of species at 50. For each case we performed 1000 simulations.

### Model fitting

We fit both LMM and PLMM to the simulated datasets with R package pez (Pearse et al. 2015). The LMM has the form

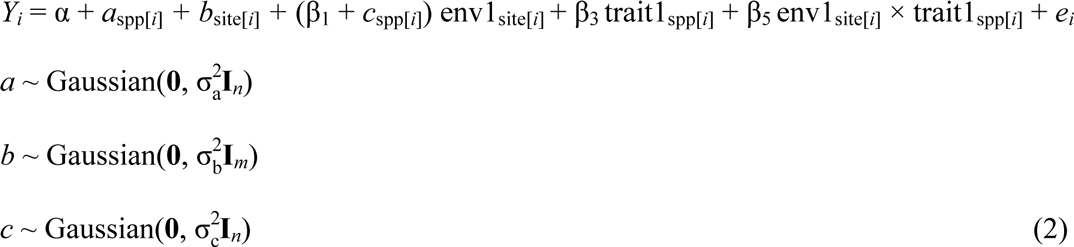

Here, we use the convention of multilevel models (Gelman & Hill, 2007), with fixed and random effects given by Greek and Latin letters, respectively. The fixed effects β_1_, β_3_, and β_5_ correspond to the same coefficients in the simulation model (equation 1). Random effect *a*_spp[*i*]_ allows different species to have different overall abundance to capture effects of the term β_4_ trait2_spp[*i*]_ in equation 1. Random effect *b*_site[*i*]_ allows different sites to have different overall abundance across all species within that site to capture effects of the term β_2_ env2_site[*i*]_ in equation 1. Finally, random effect *c*_spp[*i*]_ allows different species to have different responses to env1 to capture effects of the term β_6_ env1_site[*i*]_ × trait2_spp[*i*]_ in equation 1.

The PLMM includes all terms of equation 2, plus phylogenetic version of random terms *a*_spp[*i*]_ and *c*_spp[*i*]_:

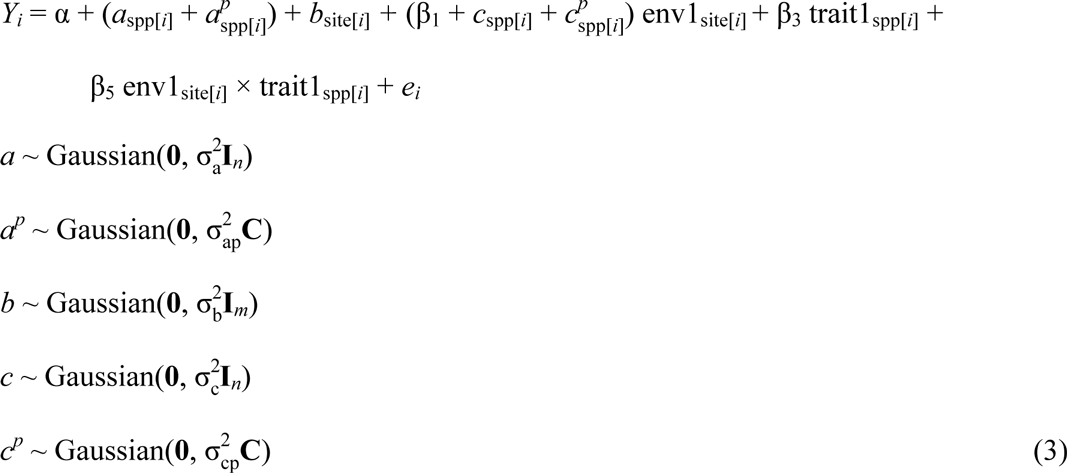

Random effect 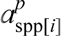 implies closely related species to have similar overall abundance; this will capture the main effects of traits in the simulations (equation 1) if trait2 has phylogenetic signal. Similarly, random effect 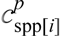 allows closely related species to have similar responses to env1, thereby capturing the interactive effect of trait2 and env1 in the simulations if trait2 has phylogenetic signal.

## Results

To compare LMMs and PLMMs, we focused on the regression coefficient β_5_ for the interaction between env1 and trait1. For each simulated dataset, we compared the accuracy of LMM and PLMM by determining the frequency with which one gave a more accurate estimate of β_5_ than the other, and also by calculating the means and standard deviations of the estimates of β_5_. We also counted the number of estimates that were scored as significant at the α = 0.05 level for both models to determine their type I errors (when the true value of β_5_ = 0) and statistical power (when the true value of β_5_ > 0).

### No phylogenetic signal in trait2

When the unmeasured trait2 did not have phylogenetic signal (trait1: **I**; trait2: **I**, and trait1: **C**; trait2: **I**), implying no phylogenetic signal in the unexplained variation in species abundances among sites, LMM and PLMM had similar estimation accuracy (Fig. 1–2), type I error rates, and power (Fig. 3). Averaged across all simulation scenarios, in roughly 50% of simulations LMM produced better estimates (closer to the true value) of β_5_ (Fig. 1). The estimators of β_5_ from LMM and PLMM had similar means and standard deviations (Fig. 2A, Fig. 2B, Fig. A1). Furthermore, LMM and PLMM had almost identical type I error rates and power across all simulation scenarios (Fig. 3). They also gave very similar estimates when β_5_ > 0 (Fig. A2). These results are explained, in part, by the fact that in about 65% of simulations across all scenarios we investigated with no phylogenetic residual variation (trait2: **I**), the estimates of both 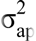 and 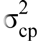 in the PLMM were zero, so the PLMM collapsed to the LMM and estimates of β_5_ were the same (± numerical accuracy in the REML optimizations).

**Figure 1.**
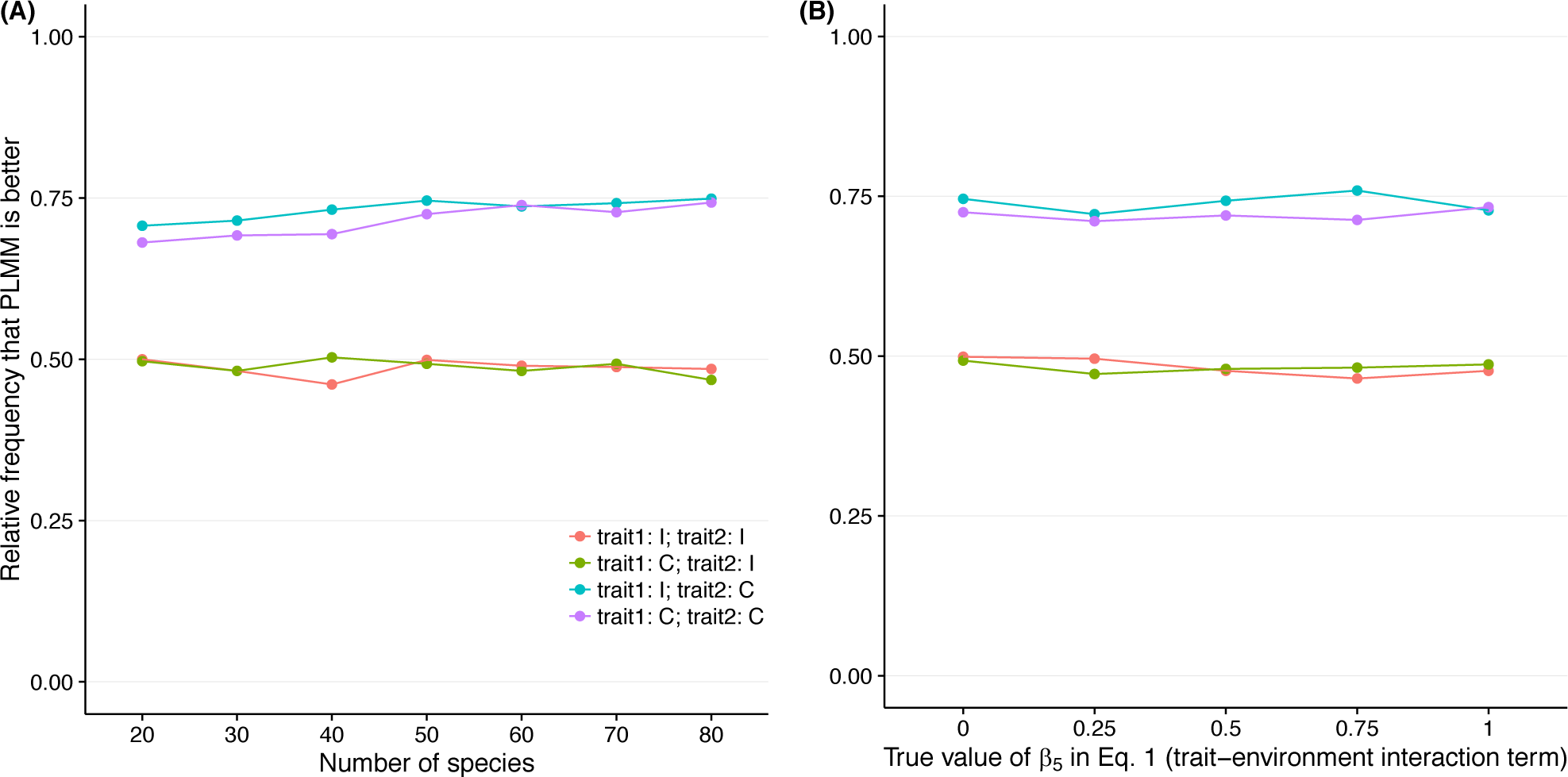
The fraction of simulations in which PLMM yielded a better estimate of β_5_ (i.e., closer to its true value) than LMM versus (A) the number of simulated species and (B) the true value of β_5_ (Eq. 1). The performance of PLMM was consistently better than LMM whenever there was phylogenetic signal in the residual variation (caused by unmeasured trait2). Abbreviations: trait1: **I** – measured trait1 does not have phylogenetic signal; trait1: **C** – measured trait1 has phylogenetic signal; trait2: **I** – unmeasured trait2 does not have phylogenetic signal; trait2: **C** – unmeasured trait2 has phylogenetic signal.

**Figure 2.**
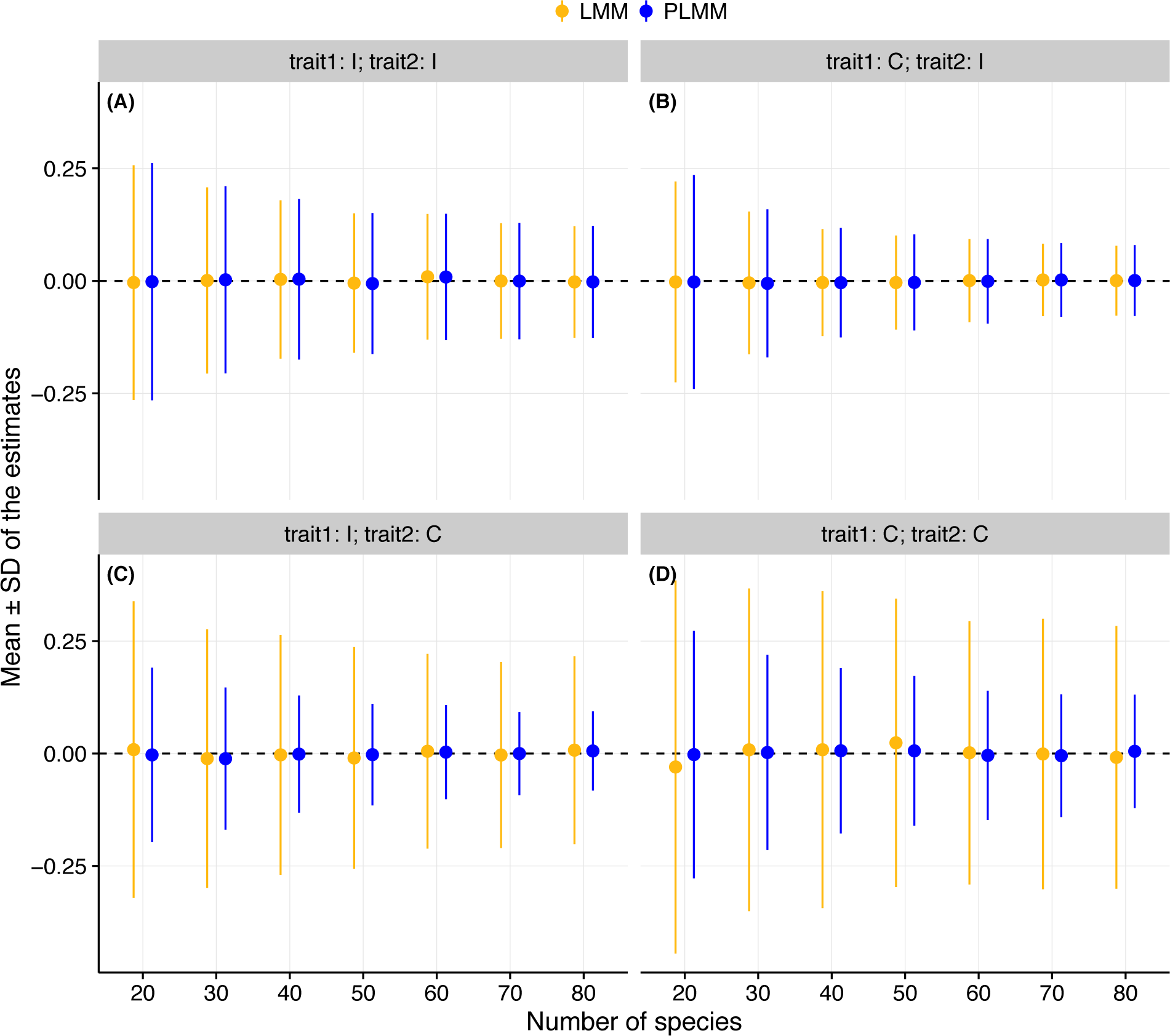
Mean (± standard deviation) of simulated estimates of β_5_ (Eq. 1) using LMM and PLMM versus the number of species in the simulations for cases (A) trait1: **I**; trait2: **I**, (B) trait1: **C**; trait2: **I**, (C) trait1: **I**; trait2: **C**, and (D) trait1: **C**; trait2: **C**. Horizontal dash lines represent the true value of the parameter. Abbreviations are as in Fig. 1.

**Figure 3.**
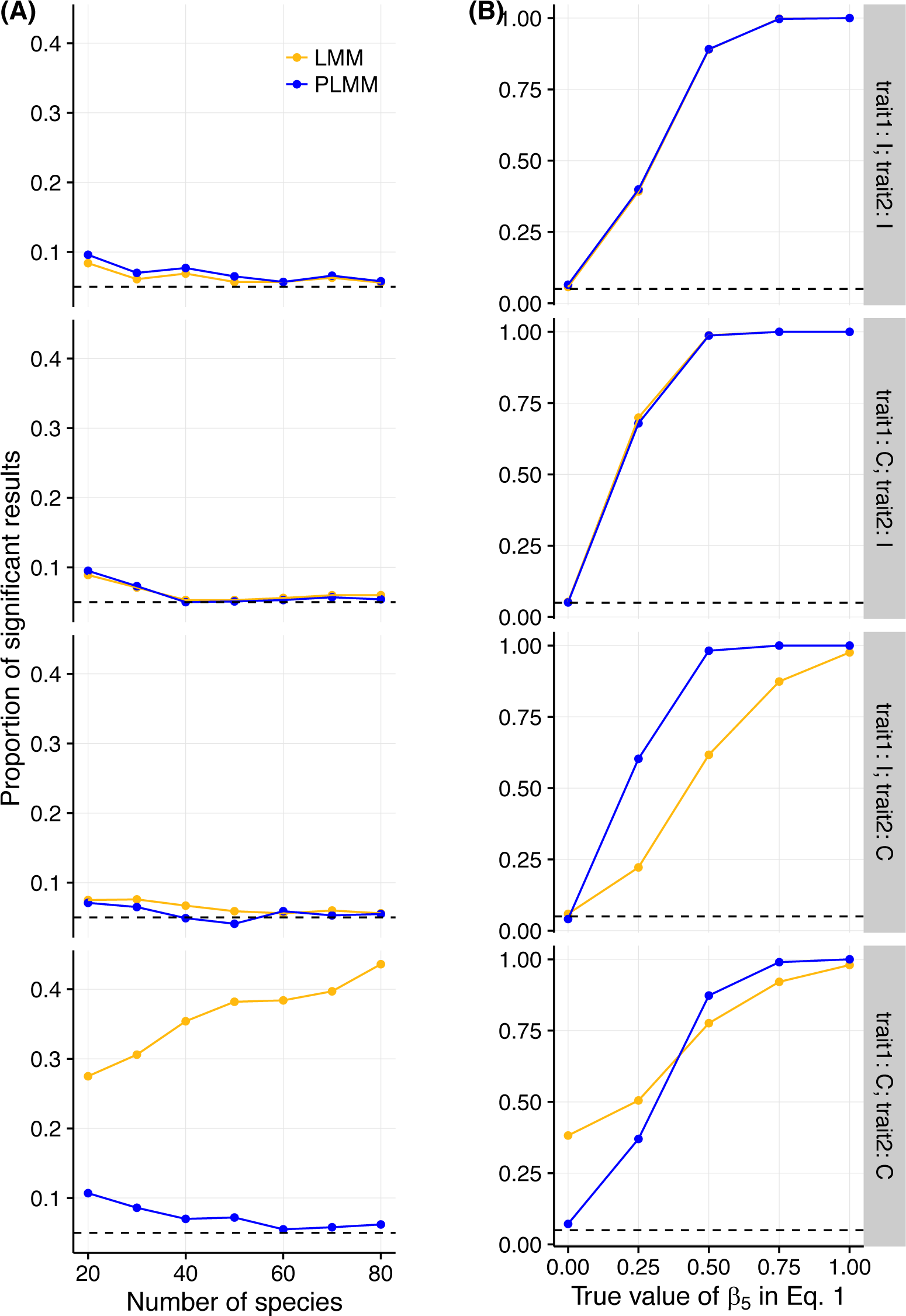
(A) Type I error rates and (B) statistical power of LMMs and PLMMs under four scenarios of simulated functional traits (abbreviations as in Fig. 1). For all tests, a significance level of α = 0.05 is used (horizontal dashed lines).

### Phylogenetic signal in trait2

When the unmeasured trait2 had phylogenetic signal (trait1: **I**; trait2: **C**, and trait1: **C**; trait2: **C**), PLMM had substantially higher estimation accuracy (Fig. 1–2), better type I error control (Fig. 3A), and higher power (Fig. 3B) than LMM. Type I error control and power were particularly poor for LMM when trait1 also had phylogenetic signal (i.e., trait1: **C**; trait2: **C**).

Averaged across all simulation conditions, in about 75% simulations PLMM produced more accurate estimates of β_5_ (Fig. 1), and the variance of the estimator of β_5_ (Fig. 2 and A1) was consistently lower than LMM. The was true regardless of the number of species, the true value of β_5_, and the status of the measured trait1 (with or without phylogenetic signal) used in simulations. In addition, for type I error control and power, LMM had particularly poor performance when the measured trait1 had phylogenetic signal (trait1: **C**; trait2: **C**). For simulations with β_5_ = 0 (Fig. 3), LMM rejected H_0_: β_5_ = 0 at the α = 0.05 level in ~25% of the datasets with 20 species, and type I error control became worse as the number of species increased. When there was no phylogenetic signal in trait1 (trait1: **I**; trait2: **C**) and type I error control was only slightly elevated, LMM had much lower power than PLMM (Fig. 3B)

We investigated further the particularly poor type I error control of LMM when there is phylogenetic signal in both the measured trait and the unexplained residual variation (trait1: **C**; trait2: **C**). Poor type I error control occurs when the estimate of the standard error of β_5_ is smaller than the true standard error. For cases both with phylogenetic signal in trait 1 (trait1: **C**; trait2: **C**) and without (trait1: **I**; trait2: **C**), we plotted the estimate of the standard error of β_5_ for each simulated dataset against the estimate of β_5_ using both LMM and PLMM (Fig. 4). For the case (trait1: **I**; trait2: **C**), the decrease in accuracy of LMM relative to PLMM is seen in the greater variance in the estimates of β_5_ (variance in the horizontal direction). Despite this increase in the variance in the estimates of β_5_, false positives (given by values to the right of the dashed line of Fig. 4) from the LMM are only slightly inflated, because the LMM estimates of the standard error of β_5_ are larger than those from PLMM. However, for the case (trait1: **C**; trait2: **C**), the decrease in accuracy of LMM relative to PLMM is not accompanied by an appropriate increase is the LMM estimates of the standard error, thereby leading to high type I error rates. In contrast to LMM, even though the variance in the estimates of β_5_ from PLMM increases when there is phylogenetic signal in trait1 (Fig. 4A vs. 4B), the estimates of the standard error also increase, leading to much better type I error control than LMM. In summary, the poor type I error control for LMM when there is phylogenetic signal in trait1 occurs because, as phylogenetic signal in trait1 increases the variance in the LMM estimates of β_5_, phylogenetic signal in trait1 decreases the LMM estimates of this variance. The decrease in power of LMM relative to PLMM for the case without phylogenetic signal in trait1 (trait1: **I**; trait2: **C**) is caused by the increase in variance in the estimator of β_5_, that is, decreased accuracy. Given the very poor type I error control for LMM for the case with phylogenetic signal in trait1 (trait1: **C**; trait2: **C**), it is inappropriate to assess power for this case.

**Figure 4.**
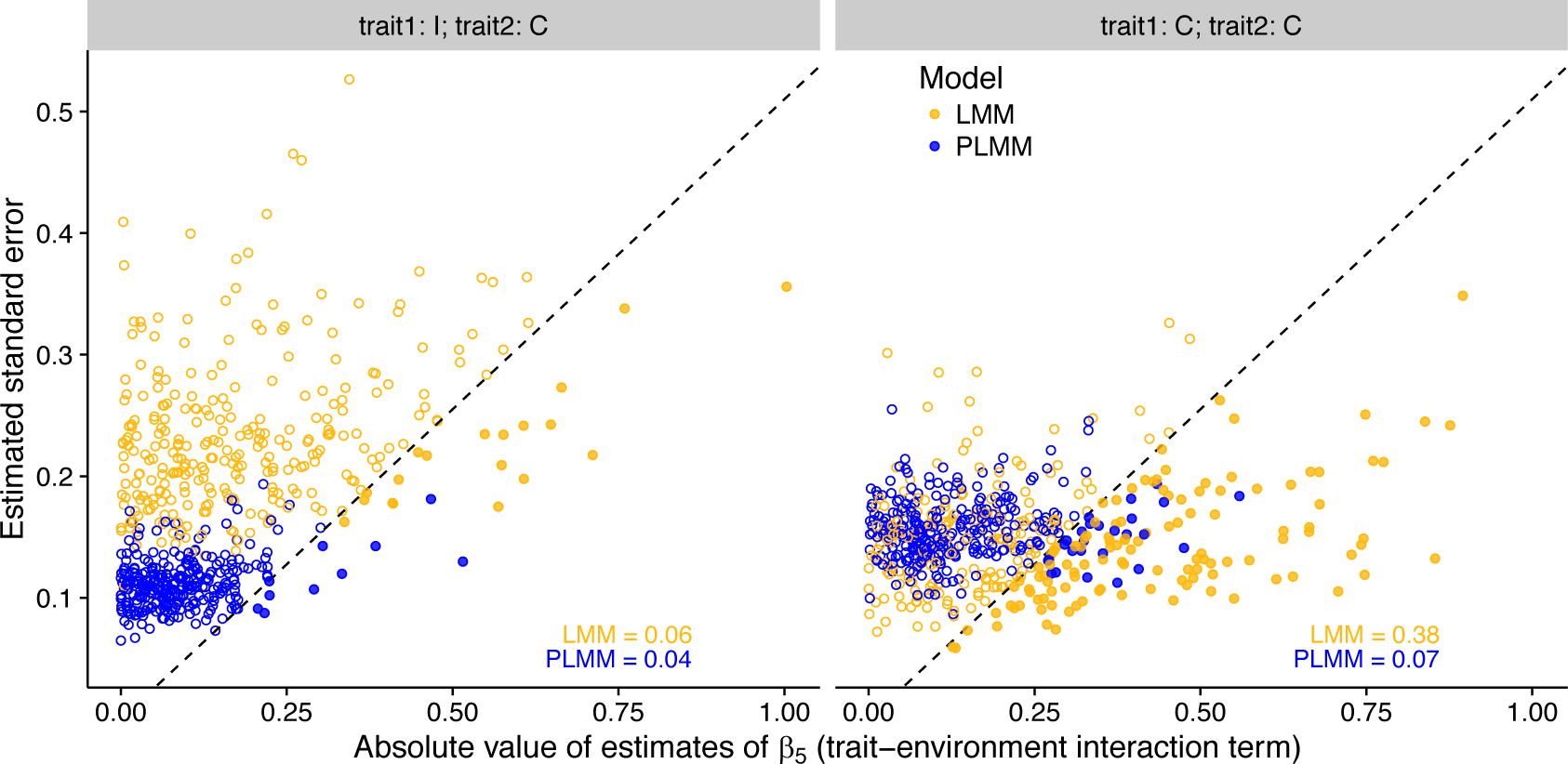
The relationships between the absolute value of the estimates of β_5_ and the estimates of the standard errors from LMM (yellow) and PLMM (blue). Solid points to the right of the dashed line would (approximately) reject the null hypothesis H_0_: β_5_ = 0 at the 0.05 α significance level. When unmeasured trait2 has phylogenetic signal but not measured trait1 (trait1: **I**; trait2: **C**), LMM estimates are more variable (horizontal axis) and have greater estimated standard errors (vertical axis) than PLMM, leading to only slightly inflated type I error control. When both trait1 and trait2 have phylogenetic signal (trait1: **C**; trait2: **C**), LMM estimates are more variable, but this is not correctly captured by increasing estimates of standard errors, leading to very high type I error rates. The dashed line has intercept of zero and slope of 1/1.96. Simulations were performed with 50 species; the fractions of simulations rejecting the null hypothesis (text in the panels) were calculated from 1000 simulations, of which only 300 are presented for clarity.

## Discussion

Our simulations have demonstrated the importance of incorporating phylogeny into the study of how species functional traits interact with the environment to affect their abundance. In simulations in which there was phylogenetic signal in the residual variation in abundances caused by an unmeasured (latent) trait, we showed that LMMs have lower accuracy, poor type I error control, and lower power than PLMMs in identifying the trait × environment interaction. The performance of LMMs was particularly poor in terms of type I error control and power when there was also phylogenetic signal in the measured trait. In contrast, PLMMs had better accuracy, generally good type I error control (except when the number of species was small), and good power.

Our results mirror the results of Revell (2010) who studied the performance of LMs and PLMs applied to regression for phylogenetic comparative data. The model he considered that most closely corresponds to our PLMM is a phylogenetic least-squares model in which Pagel’s λ branch-length transform is used. Pagel’s λ transformation can be constructed by adding a phylogenetic and a non-phylogenetic covariance matrix with λ scaling between them (i.e., (1 –λ)**I** + λ**C**). In our PLMM (Eq. 3), covariance terms are similarly combined; for example, the covariance for species-specific slopes across environmental variable 1 is 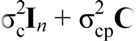. Revell (2010) found that PLMs outperformed LMs when there was phylogenetic signal in the residual variation, with the performance of LMs particularly poor when there was also phylogenetic signal in the independent variable. Thus, we found similar results in the more-complex problem of identifying trait × environment interactions in community data.

The better performance of PLMMs over LMMs is not surprising on theoretical grounds. For the special, hypothetical case in which the variance parameters 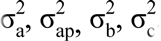, and 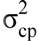 are known, the PLMM in equation 3 will be the minimum variance estimator of the regression coefficients (fixed effects), including the trait × environment interaction β_5_; this is a consequence of the Cramer-Rao Theorem applied to Generalized Least Squares (GLS) models (Judge et al. 1985). This explains why PLMMs provide more accurate estimates of β_5_ than LMMs, and the increase in accuracy explains the increase in power of PLMMs relative to LMMs.

A particular warning derived from our simulations is the poor type I error control for LMMs when there is phylogenetic signal in both the residual variation and in the independent variable. When there is also phylogenetic signal in the measured trait1, the variance in the estimates of β_5_ greatly increases. Nonetheless, the LMM estimates of the standard error of β_5_ do not increase as they should, leading to false rejections of the null hypothesis that β_5_ = 0. Because PGLMMs are close to the minimum variance estimators of β_5_, the variance in its estimates of β_5_ does not increase as much as LMMs when there is phylogenetic signal in the independent variable, and what increase occurs is correctly given by the estimates standard errors of β_5_; thus, there is generally good type I error control.

When the number of species is small (<60), however, PLMM had inflated type I error rates; for simulations with 20 species and phylogenetic signal in both independent variable (measured trait1) and residual variation (unmeasured trait2), the null hypothesis H_0_: β_5_ = 0 was rejected in 10% of the datasets at the a significance level of 0.05. In analyses with small numbers of species and P-values computed from the data that are close to the significance level selected by the researcher, we suggest using parametric bootstrapping. This can be performed by estimating parameters from the data under H_0_: β_5_ = 0 (i.e., without the trait × environment interaction), simulating a large number (e.g., 2000) datasets with these parameter values, fitting each dataset with the full model (i.e., with the trait × environment interaction), and for each dataset recording the Z-score of the estimate of β_5_. The bootstrap approximate P-value of β_5_ under an a significance level of 0.05 is then given by the proportion of bootstrap Z-scores whose absolute values exceed the absolute value of the Z-score from the observed data. Code for performing this bootstrap is provided in Appendix S1.

Our analyses have been confined to abundance as a continuous dependent variable. Presence/absence (incidence) community data can also be analysed with phylogenetic information using PGLMM (Ives and Helmus 2011), and results will likely be similar. We did not pursue this here, however, because the computational burden of PGLMMs with existing software makes simulation studies difficult. Nonetheless, if tests of the existence of relationships (i.e., testing H_0_: β_5_ = 0) are all that is needed, applying PLMMs to binary data generally provides good type I error control, although at the expense of some power (Ives 2015; Warton et al. 2016).

Even when there was no phylogenetic signal in the residual variation, PLMMs performed as well as LMMs. In part, this is because, when PLMMs detected no phylogenetic signal in the residual variation, they give the same results as the corresponding LMMs (although their AIC values are still penalized by the variance term that equals zero). The fact that PLMMs often collapse exactly to LMMs as a special case suggests that PLMMs should be always used in analyses of trait × environment interactions, since there is no cost in the absence of phylogenetic signal and considerable benefits when there is (which is likely).

## Acknowledgements

This work was supported by NSF/NASA-DEB-Dimensions 1240804 (to ARI) and NSF DEB-Dimensions 1046355.

## Data accessibility

No data were used in this article.

## Supporting information

Appendix S1: R scripts used for the simulations and analyses (in the Rcode folder). It is time consuming to re-run all simulations and model fittings. Therefore, we also include all model results for fixed effects (as rds.zip and rds2.zip files). (We cannot upload these two files through manuscript central as they include >50 files; one can download them from here: https://uwmadison.box.com/s/l1nnsmqlymg53xyxdarfksjrtegvuzci)

**Figure A1.**
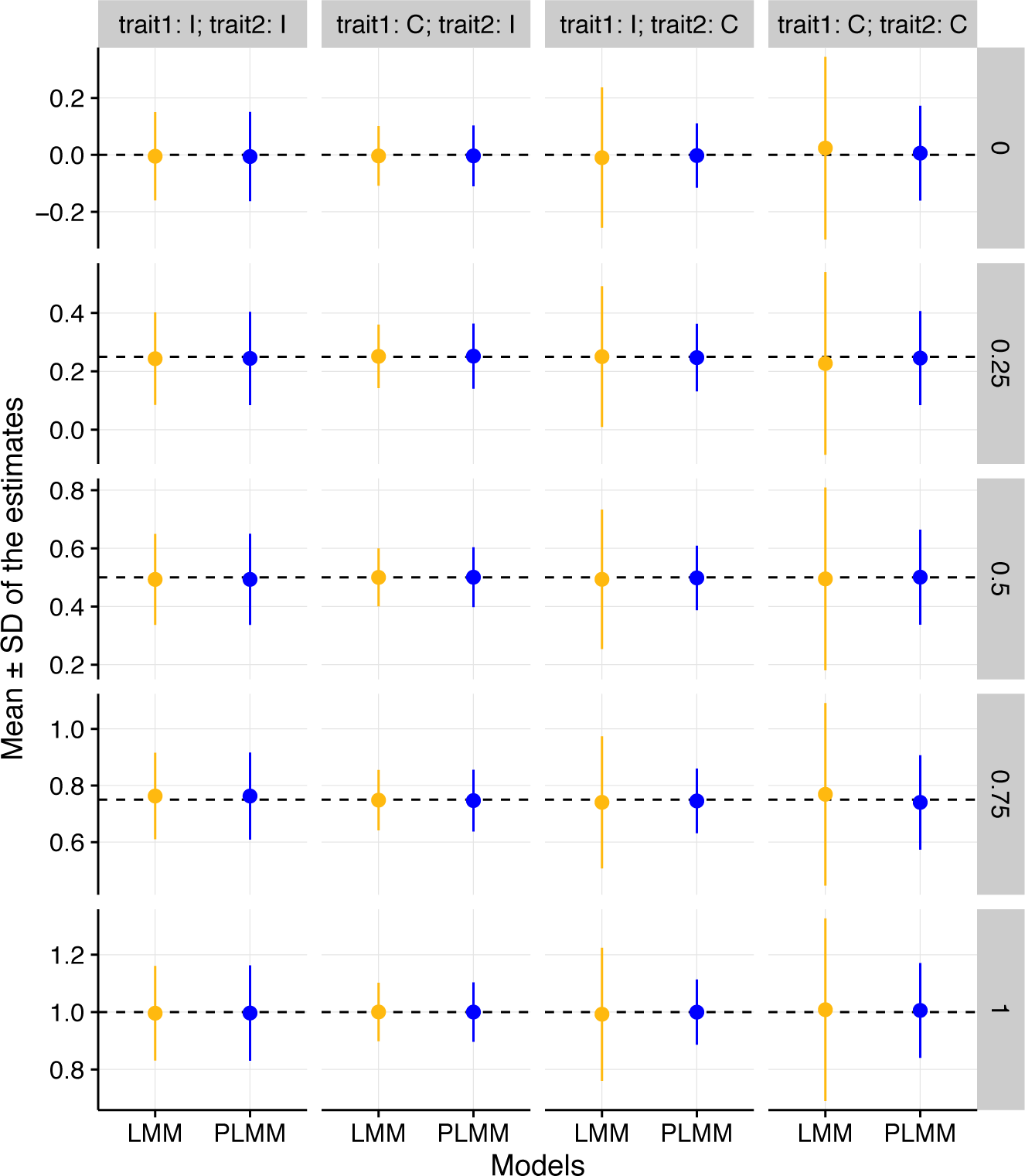
Mean (± standard deviation) of estimates of β_5_ (Eq. 1) using LMM and PLMM varying the true value of β_5_ (0, 0.25, 0.5, 0.75, and 1). Horizontal dash lines represent the true value of the parameter. Abbreviations are as in Fig. 1.

**Figure A2.**
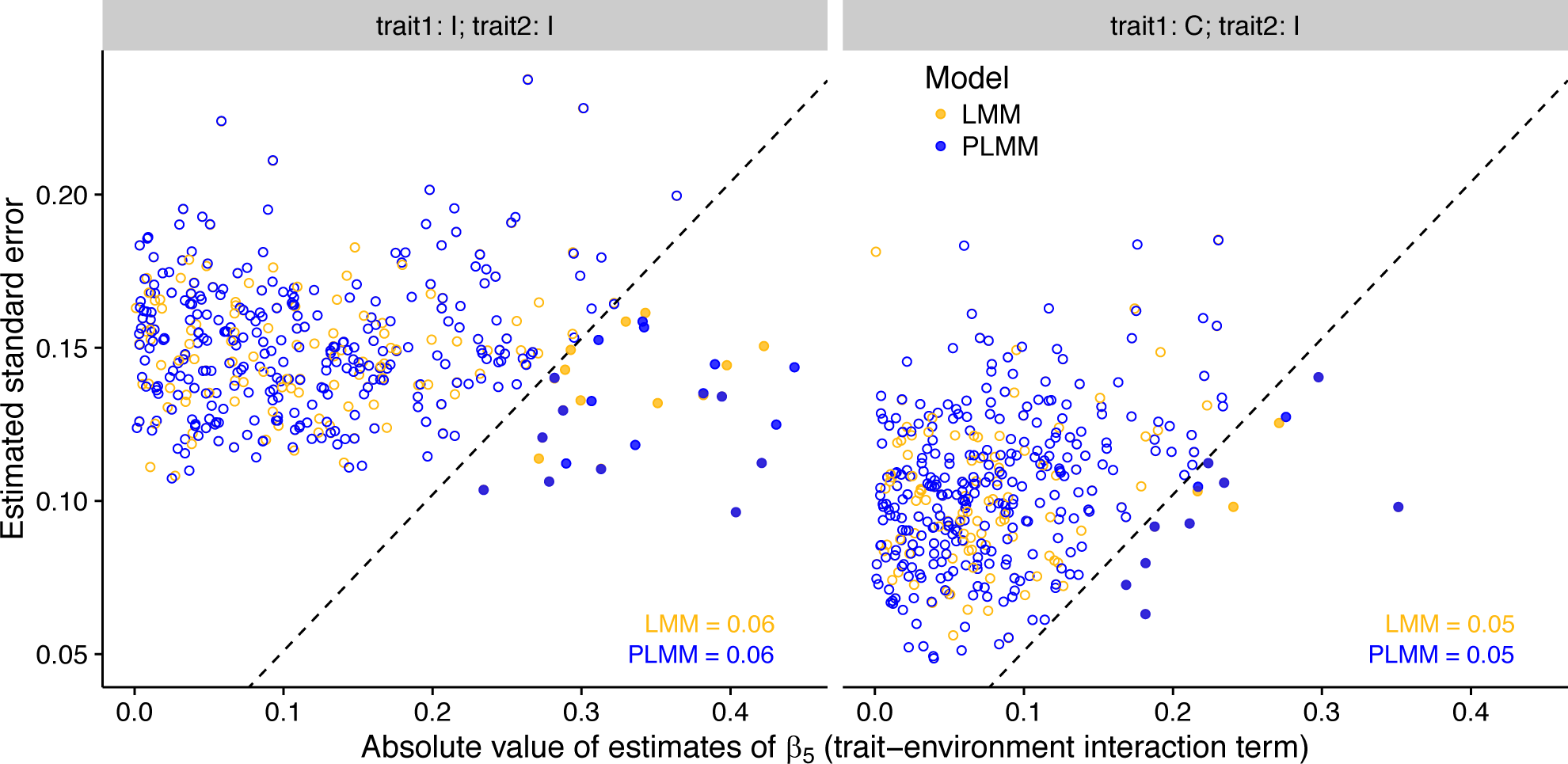
The relationships between the absolute value of the estimates of β_5_ and the estimates of the standard errors for LMM (yellow) and PLMM (blue). Solid points to the right of the dashed line would reject the null hypothesis H_0_: β_5_ = 0. When neither trait1 nor trait2 has phylogenetic signal (trait1: **I**; trait2: **I**), LMM and PLMM estimates have similar variability (horizontal axis) and similar estimated standard errors (vertical axis). When trait1 has phylogenetic signal (trait1: **C**; trait2: **I**), both LMM and PLMM estimates become less variable, which is correctly captured by decreasing estimates of standard errors, leading to appropriate type I error rates. The dashed line has intercept of zero and slope of 1/1.96. Simulations were performed with 50 species; the fractions of simulations rejecting the null hypothesis (text in the panels) were calculated from 1000 simulations, of which only 300 are presented for clarity.

## References

Blomberg, S. P., Garland, T., & Ives, A.R. (2003). Testing for phylogenetic signal in comparative data: behavioral traits are more labile. Evolution, 57:717-745.

Bolker, B.M., Brooks, M.E., Clark, C.J., Geange, S.W., Poulsen, J.R., Stevens, M.H.H. & White, J.-S.S. (2009). Generalized linear mixed models: A practical guide for ecology and evolution. Trends in Ecology & Evolution, 24, 127–135.

Brown, A.M., Warton, D.I., Andrew, N.R., Binns, M., Cassis, G. & Gibb, H. (2014). The fourth-corner solution - using predictive models to understand how species traits interact with the environment. Methods in Ecology and Evolution, 5, 344–352.

Cavender-Bares, J., Kozak, K.H., Fine, P.V.A. & Kembel, S.W. (2009). The merging of community ecology and phylogenetic biology. Ecology Letters, 12, 693–715.

Dray, S. & Legendre, P. (2008). Testing the species traits-environment relationships: The fourth-corner problem revisited. Ecology, 89, 3400–3412.

Felsenstein, J. (1985). Phylogenies and the Comparative Method. The American Naturalist, 125, 1–15.

Garamszegi, L.Z. (2014). Modern phylogenetic comparative methods and their application in evolutionary biology. Concepts and Practice. London, UK: Springer.

Garland, T., Bennett, A.F. & Rezende, E.L. (2005). Phylogenetic approaches in comparative physiology. Journal of experimental Biology, 208, 3015–3035.

Gelman, A. & Hill, J. (2007). Data analysis using regression and multilevel/hierarchical models. Cambridge University Press.

Grafen, A. (1989). The phylogenetic regression. Philosophical Transactions of the Royal Society of London. Series B, Biological Sciences, 326, 119–157.

Harmon, L., Weir, J., Brock, C., Glor, R. & Challenger, W. (2008). GEIGER: Investigating evolutionary radiations. Bioinformatics, 24, 129–131.

Harvey, P.H., Pagel, M.D. & others. (1991). The comparative method in evolutionary biology. Oxford university press Oxford.

Ives, A.R. (2015). For testing the significance of regression coefficients, go ahead and log-transform count data. Methods in Ecology and Evolution, 6, 828–835.

Ives, A.R. & Helmus, M.R. (2011). Generalized linear mixed models for phylogenetic analyses of community structure. Ecological Monographs, 81, 511–525.

Jackson, M.M., Turner, M.G., Pearson, S.M. & Ives, A.R. (2012). Seeing the forest and the trees: Multilevel models reveal both species and community patterns. Ecosphere, 3, art79.

Jamil, T., Ozinga, W.A., Kleyer, M. & ter Braak, C.J. (2013). Selecting traits that explain speciesenvironment relationships: A generalized linear mixed model approach. Journal of Vegetation Science, 24, 988–1000.

Judge, G.G., Griffiths, W., Hill, R.C., Lutkepohl, H. & Lee, T.C. (1985). Introduction to the theory and practice of econometrics.

Legendre P., M.L.H.-V., René Galzin. (1997). Relating behavior to habitat: Solutions to the fourth-corner problem. Ecology, 78, 547–562.

Martins, E.P. & Hansen, T.F. (1997). Phylogenies and the Comparative Method: A General Approach to Incorporating Phylogenetic Information into the Analysis of Interspecific Data. The American Naturalist, 149, 646–667.

McGill, B.J., Enquist, B.J., Weiher, E. & Westoby, M. (2006). Rebuilding community ecology from functional traits. Trends in Ecology & Evolution, 21, 178–185.

Ovaskainen, O., De Knegt, H.J. & Mar Delgado, M. del. (2016). Quantitative ecology and evolutionary biology: Integrating models with data. Oxford University Press.

Paradis, E. (2011). Analysis of phylogenetics and evolution with r. Springer Science & Business Media.

Pearse, W.D., Cadotte, M.W., Cavender-Bares, J., Ives, A.R., Tucker, C.M., Walker, S.C., & Helmus, M.R. (2015). pez: phylogenetics for the environmental sciences. Bioinformatics, 31, 2888–2890.

Pollock, L.J., Morris, W.K. & Vesk, P.A. (2012). The role of functional traits in species distributions revealed through a hierarchical model. Ecography, 35, 716–725.

R Core Team. (2015). R: A Language and Environment for Statistical Computing.

Revell, L.J. (2010). Phylogenetic signal and linear regression on species data. Methods in Ecology and Evolution, 1, 319–329.

Revell, L.J. (2012). Phytools: An r package for phylogenetic comparative biology (and other things). Methods in Ecology and Evolution, 3, 217–223.

Soudzilovskaia, N.A., Elumeeva, T.G., Onipchenko, V.G., Shidakov, I.I., Salpagarova, F.S., Khubiev, A.B., Tekeev, D.K. & Cornelissen, J.H.C. (2013). Functional traits predict relationship between plant abundance dynamic and long-term climate warming. Proceedings of the National Academy of Sciences, 110, 18180–18184.

Warton, D.I., Foster, S.D., De’ath, G., Stoklosa, J. & Dunstan, P.K. (2014). Model-based thinking for community ecology. Plant Ecology, 1–14.

Warton, D.I., Lyons, M., Stoklosa, J. & Ives, A.R. (2016). Three points to consider when choosing a LM or GLM test for count data. Methods in Ecology and Evolution, 7, 882–890.

Webb, C.O., Ackerly, D.D., McPeek, M.A. & Donoghue, M.J. (2002). Phylogenies and community ecology. Annual Review of Ecology and Systematics, 33, 475–505.

Westoby, M. & Wright, I.J. (2006). Land-plant ecology on the basis of functional traits. Trends in Ecology & Evolution, 21, 261–268.

